# Unlocking cross-modal interplay of single-cell and spatial joint profiling with CellMATE

**DOI:** 10.1101/2024.09.06.610031

**Authors:** Qi Wang, Bolei Zhang, Luyu Gong, Yue Guo, Erguang Li, Jingping Yang

## Abstract

A key advantage of single-cell multimodal joint profiling is the modality interplay, which is essential for deciphering the cell fate. However, while current analytical methods can leverage the additive benefits, they fall short to explore the synergistic insights of joint profiling, thereby diminishing the advantage of joint profiling. Here, we introduce CellMATE, a <u>M</u>ulti-head <u>A</u>dversarial <u>T</u>raining-based <u>E</u>arly-integration approach specifically developed for multimodal joint profiling. CellMATE can capture both additive and synergistic benefits inherent in joint profiling through auto-learning of multimodal distributions and simultaneously represents all features into a unified latent space. Through extensive evaluation across diverse joint profiling scenarios, CellMATE demonstrated its superiority in ensuring utility of cross-modal properties, uncovering cellular heterogeneity and plasticity, and delineating differentiation trajectories. CellMATE uniquely unlocks the full potential of joint profiling to elucidate the dynamic nature of cells during critical processes as differentiation, development and diseases.

**Graphical abstracts:** 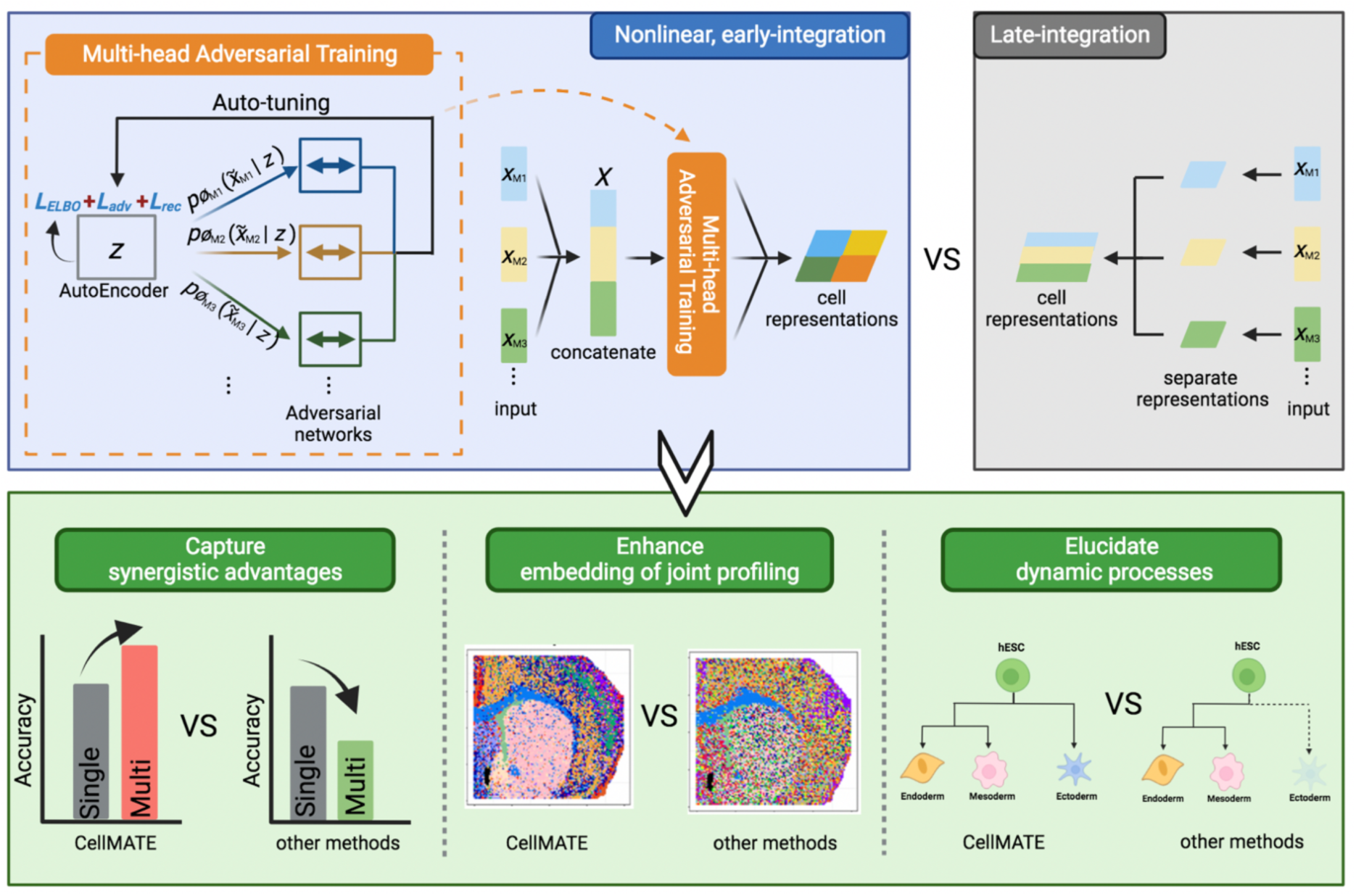

## INTRODUCTION

Cell fates are jointly determined by a spectrum of molecular characteristics, including gene expression, chromatin accessibility and other properties. The interplay across these molecular modalities is especially crucial for cells undergoing dynamic processes such as differentiation, development and diseases[1–3]. Recent advancements in experimental techniques have enabled single-cell multimodal (sc-multimodal) joint profiling and generated valuable paired sc-multimodal data that could shed light on cellular heterogeneity and plasticity[4–7]. However, while current analytical methods can leverage the additive benefits in paired sc-multimodal data[5, 8], they fall short in capturing the synergistic insights across modalities. This gap underscores a pressing need for analytical approaches that can capture the synergistic benefits of paired multimodal data to harness the full potential of joint profiling.

Current analytical methods cannot harness synergistic benefits due to the late integration or linear modelling [9–15]. Late integration methods like WNN[9], segregate modalities into distinct latent spaces before synthesis, and assume that cellular states defined across modalities do not conflict[5, 9]. However, this assumption often neglects the subtle yet crucial signals within individual modalities during joint analysis, and fail to capture the cross-modal interplay inherent in joint profiling[16]. This is particularly detrimental for understanding the dynamic nature of cells as evidenced by lineage priming during differentiation[17]. Alternatively, early integration methods with linear modelling like MOFA+[11], though not constrained by aforementioned limitations, struggle with the intrinsic non-linearity of sc-multimodal data[5, 8]. Therefore, enabling early integration and accounting for non-linearities are essential for capturing the synergistic advantages in paired multimodal data.

Deep generative frameworks, which have recently been widely applied on single cell data[14, 18, 19], can learn complex non-linearities[5]. However, early integrating diverse distributions and noises from various modalities into a singular such framework remains a challenge[20, 21]. Furthermore, existing deep generative frameworks for single-cell data typically use off-the-shelf parametric modelling, which in practice may not well model the true distribution and noise of sc-multimodal data[21, 22]. Moreover, the rapid evolving joint profiling experimental techniques are continuously expanding the types and numbers of modalities simultaneously profiled within single cells[2, 20, 23–27], complicating the analytic task. Hence, achieving precise interpretation of paired multimodal data necessitates a strategy adept at accurately and simultaneously addressing the diverse distributions and noises across modalities.

To tackle these challenges, we introduce CellMATE, a <u>M</u>ulti-head <u>A</u>dversarial <u>T</u>raining enabled <u>E</u>arly-integration approach, specifically designed to harness the full potential of paired sc-multimodal data. CellMATE’s architecture uniquely enables auto-learning of varied distribution and noise attributes directly from real paired sc-multimodal data and simultaneously represents all features into a unified latent space. By enabling early and nonlinear integration of sc-multimodal data, CellMATE adeptly captures both the additive and synergistic advantages of joint profiling. Our comprehensive benchmarks demonstrate that CellMATE is robust across diverse paired sc-multimodal scenarios, showcasing its unparalleled capability to elucidate synergistic strength even amidst modal discrepancies. CellMATE thus harness the full potential of joint profiling, providing insight into critical and dynamic processes.

## MATERIAL AND METHODS

### Model Architecture of CellMATE

The main framework of CellMATE is a deep generative framework uniquely utilizing a multi-head adversarial training module. Given multimodal data 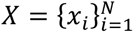, where N is the number of cells, we represent each cell *x_i_* ∈ *X* as a concatenation of features from *M* modalities, i.e., 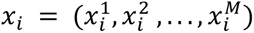. A modal-free encoder *q_θ_* (*z_i_* |*x_i_*) takes the whole input multimodal data *x_i_* without distinguishing their modality differences and encodes features from all modalities into a modality-independent latent representation *z_i_*, where *θ* is the parameter of the encoder. A standard Gaussian distribution with mean *μ* and log-variance *log σ* is used for *z*, while introducing the necessary randomness: *z_i_* = *μ_i_* + *ϵ σ_i_*, where *ϵ* ∼ 𝒩(0, *I*). The latent representations are then projected back into the original high-dimensional input space using modal-specific adversarial networks. The modal-specific adversarial networks consist of a multi-head decoder and multiple modal-specific discriminators. To enable the recovery of accurate modality-specific features from *z* and to capture the true and modality-specific data distribution and noise instead of relying on off-the-shelf parametric modeling, the decoder network *p_ϕ_*(**x*_i_*|*z_i_*), which is parameterized by *ϕ*, uses a multi-head fully connected layer with each head responsible for generating features of a corresponding modality, and multiple modal-specific discriminators *D*_*ψ*_*^j^* are used, with the *j*th modality is parameterized by *ψ*_*j*_. To save time, the adversarial training starts with negative binomial (NB) distribution for scRNA and scADT and zero-inflated negative binomial (ZINB) distribution for scATAC and scChIP, and is then tuned by minimizing the f-divergence between the true and reconstructed data. The true and modality-specific data distribution and noise can then be effectively recovered by the adversarial training, and the accurate joint latent representation can be inferred by maximizing the similarity of the regenerated data and the input data.

Training CellMATE is based on maximizing the log-likelihood of the observed features, which can be formulated as:

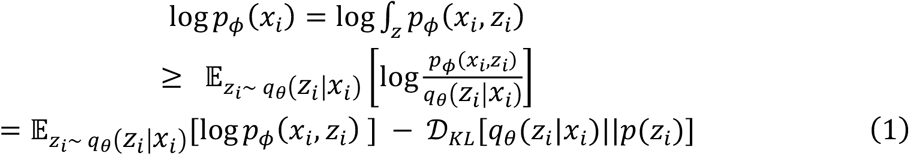

where *p*(*z_i_*) is assumed to be a normal distribution with 0 mean and unit variance. Then according to the evidence lower bound (ELBO), the cost function is formulated as:

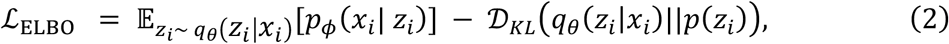

The encoder and decoder network are also optimized by considering the loss of adversarial training, which can be formulated as:

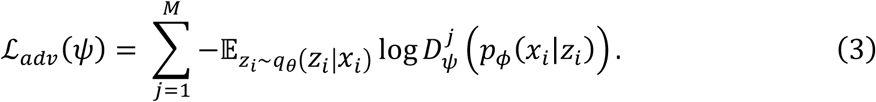

However, since the network has difficulty mapping the generated samples to the latent space, and as a result of that, the ELBO lags behind or even increases during the training process[28]. In order to solve this problem, we incorporate an additional recovery loss, which uses *l*_1_ distance to measure the distance between *z* and recovered *q_θ_*(*p_ϕ_*(⋅ |*z*)). The loss can be formulated as:

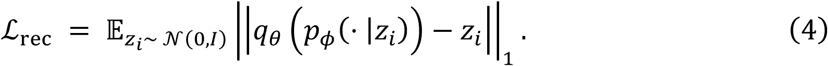

In summary, we minimize the cost function of the encoder and the decoder network as a weighted sum of the above functions:

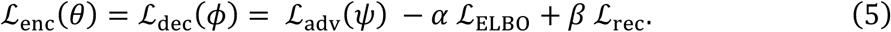

where *α* and *β* are non-negative hyperparameters. In this study, we set *α* = 0.1, *β* = 1.

For the discriminator in the adversarial network, there are 3 sources of input: the real data *x*, the regenerated data *x* from encoded latent *z*, and random prior *x̅_i_*^j^ from normal distribution 𝒩(0, *I*). For each modality *j*, the discriminator should output 1 for *x_i_^j^*, 0 for *x_i_^j^*, and 0 for *x̅_i_^j^*, thus the cost function of *D*_*ψ*_*^j^* (⋅) can then be formulated as cross-entropy function:

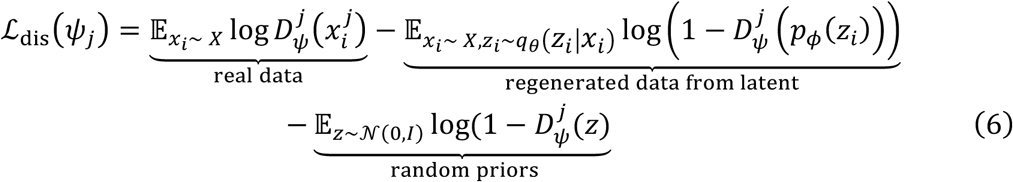

The overall network structure of CellMATE is as follows. The modal-free encoder network is constructed with a multi-layer neural network, including a fully connected layer for mean *μ* and variance *σ* of the latent representation *z*, a 1-dimensional batch normalization layer and a ReLU layer. The hidden layers of the multi-head decoder network consist of a multi-head fully connected layer for generating features of each modality, a 1-dimensional batch normalization layer and a ReLU layer. Each discriminator in the adversarial network has 3 composite layers, with each composite layer consisting one fully connected layer, one 1-dimensional batch normalization layer, one leaky ReLU layer and one dropout layer. The output of the discriminator is a fully connected layer with Sigmoid to output the probability of being real or fake distributions.

### Model Training

For the parameters updating, we use Adam as the optimizer for updating the gradients. By default, the learning rate is set to 0.0001, the weight decay is set to 0.000001, the batch size is set to 128 and the training epoch is set to 1000. After optimizing the neural networks, we adopt clustering algorithm with the latent representation z. In this paper, we use the Louvain clustering algorithm which has been used for community detection to extract communities from large networks.

### Benchmarking and evaluation

#### Data preprocessing

We used a total of eight datasets for benchmarking. For feature selection in each dataset, low-quality features and lowly-variable features with abnormally high coverage were filtered out. If not emphasized specifically, low-quality features were defined as features detected in fewer than 1% of the cells and lowly-variable features were defined as features detected in more than 95% of the cells.

The PBMCs (10X) dataset was downloaded from the 10X genomics website (https://www.10xgenomics.com/resources/datasets/pbmc-from-a-healthy-donor-granulocytes-removed-through-cell-sorting-3-k-1-standard-1-0-0). The kidney (sci-CAR) dataset was downloaded from the Gene Expression Omnibus (GEO) repository under the accession numbers GSM3271044. Low-quality genes were defined as genes with no coverage, and lowly-variable features were not defined for this dataset. The spatial RNA+ATAC dataset was obtained from UCSC Cell and Genome Browser (https://brain-spatialomics.cells.ucsc.edu). Lowly-variable genes were defined as genes detected in more than 50% of cells. The MulTI-Tag dataset was downloaded from the Zenodo repository (https://doi.org/10.5281/ zenodo.6636675), and feature selection was done according to the provided codes in the original manuscript. The nanobody-based scCUT&Tag dataset was downloaded from GSE198467. The ASAP-seq, CITE-seq and DOGMA-seq dataset was downloaded from GSE156478. For the ASAP-seq dataset, peak-by-cell matrix was generated from the fragments file using the ArchR package. Low-quality proteins were defined as proteins with no coverage, and lowly-variable proteins were defined as proteins detected in 100% of the cells.

For methods as Cobolt, MultiVI, scAI, scMM, scDEC that require transcriptome or ADT, GeneScore matrices were generated from other modalities in datasets lacking transcriptome or ADT using the ArchR package. For scRNA, the raw count matrix is directly used as input, for scATAC and scChIP, the raw counts are binarized and then used as input, and for scADT, the data is normalized using the “NormalizeData” function with the following parameters “normalization.method = ’CLR’, margin = 2” in ‘Seurat’ R package.

#### Cell type annotation

For the kidney (sci-CAR) dataset, cell type labels were defined according to the original manuscript[29]. For the MulTI-Tag dataset, cell-type labels were downloaded from the Zenodo repository (https://doi.org/10.5281/ zenodo.6636675). For the nanobody-based scCUT&Tag dataset, cell-type labels were defined if labels from two out of three characteristics (ATAC, H3K27me3 and H3K27ac) provided were consistent, otherwise, cell-type labels from ATAC were used instead. For datasets without cell-type labels (the PBMCs (10X), the CITE-seq and the DOGMA-seq datasets), the ground truth cell-type labels were inferred according to the Seurat “Multimodal reference mapping” tutorial with annotated PBMC reference[30] provided in the vignettes. Since scRNA-seq was absent in ASAP-seq dataset, the cell-type labels were inferred according to the Seurat “Integrating scRNA-seq and scATAC-seq data” tutorial with Seurat annotated PBMC reference.

#### Settings used in competing methods

WNN[9] (Seurat V4), MOFA+[11], scAI[10], Cobolt[13], MultiVI[14], scMM[12] and scDEC[15] were used as competing methods. For each method, if not emphasized specifically, we all used the default parameter settings as recommended in the tutorial or parameter settings used in the original manuscript.

##### WNN

WNN was executed using an R (4.1.0) package ‘Seurat’ (v 4.1.1) according to the website tutorial (https://satijalab.org/seurat). The latent dimension of each modality was fixed to 24 and then we calculated the closest neighbours for each cell based on a weighted combination of each modality using the “FindMultiModalNeighbors” function. Louvain clustering was used in the “FindClusters” step.

##### MOFA+

MOFA+ was executed using R (4.1.0) package ‘MOFA2’ (v. 1.4.0) according to the website tutorial (https://github.com/bioFAM/MOFA). Variable features were identified for each modality using the “FindVariableFeatures” function. To train the mofa model, “create_mofa” function was used and the number of factors was set to 24.

##### scAI

scAI was executed as implemented from its source code using the R (4.1.0) package ‘scAI’ (v. 1.0.0). To run the scAI model, we used the “run_scAI” function with the following parameters “K = 24, nrun = 1, do.fast = T”. Clusters were then identified using the “identifyClusters” function with default parameters.

##### Cobolt

Cobolt (v. 1.0.1) was executed according to the tutorial provided on github (https://github.com/epurdom/cobolt_manuscript). Multiomic dataset was created using the “MultiomicDataset.from_singledata” function. The n_latent parameter was set to 24 in the training step. For the learning rate parameter, 0.005 was applied for most datasets and 0.001 was applied for the MulTI-Tag dataset according to the error recommendations.

##### MultiVI

MultiVI was conducted using the scvi (v. 1.0.2) python package according to the authors’ API functions (scvi-tools.org) with parameters used in the provided code. Data was transformed into an AnnData object using the “read_10x_multiome” function and trained using the “train” function with default parameters. The latent space was obtained from the trained model using the “get_latent_representation” function.

##### scMM

scMM was executed according to the tutorial provided on github (https://github.com/kodaim1115/scMM) and the latent_dim parameter was set to 24.

##### scDEC

scDEC was executed according to the tutorial provided on github (https://github.com/kimmo1019/scDEC) with the following parameters “--dy 24 --no_label --mode 3 --train True”. During the model evaluation step, the newest epoch was used to generate the latent feature matrix and the inferred cluster label.

#### Evaluation metrics

To evaluate the performance of the metrics, we mainly compare the following metrics: Adjusted Rand Index (ARI), Unsupervised Clustering Accuracy (ACC), Local inverse Simpsons Index (LISI) and Over-correction (OC) score.

##### ARI

ARI was calculated by “adjusted_rand_score” function in the ‘sklearn.metrics’ package, which is a similarity score between two clustering assignments. ARI is independent of the number of clusters. The value is 0 for random prediction and 1 for perfect matching.

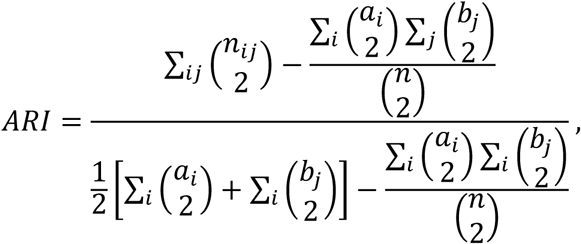

where *n_ij_*, *a_i_* and *b_j_* are values from the contingency table.

##### ACC

ACC was calculated by comparing the predicted labels and true cell-type labels using the following formula:

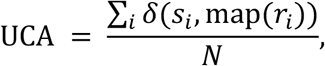

where *r*_%_ is the predicted label and *s_i_* is the true cell-type label; *δ*(⋅) is an indication function which is 1 if *s_i_* equals map(*r_i_*) ; map(⋅) is a optimal rearrange method which can be implemented by Hungarian Algorithm.

##### LISI

LISI values were calculated for each cell using the “compute_lisi” function in the ‘lisi’ (v. 1.0) R package. 2D-embeddings and cell-type labels were used as input. The averaged LISI values across all cells were calculated for each method.

##### OC score

OC score was calculated as defined in previous study[31]. Briefly, the percentage of the k-nearest neighbouring cells with distinct cell-type labels was calculated for each cell. k was set to 100 as original study defined.

### Marker feature analysis

To identify marker features in different subgroups of CD14 Monocytes, “FindMarkers” function in the ‘Seurat’ package was used. Features with “avg_log2FC > 0, p_val_adj < 0.01” were defined as subgroup marker features.

### Trajectory analysis

Trajectory inference of the differentiated hESCs was performed using ‘slingshot’ (v. 2.5.2) R package. UMAP embeddings from each method were provided as input. To identify features of H3K36me3, H3K27me3 or H3K4me1 histone modification signals significantly changed along the ectoderm trajectory, “differentialGeneTest” function in the ‘Monocle’ (v. 2.14.0) R package was used. Top 100 genes or peaks that most significantly changed along the trajectory were shown in the heatmap plot. Smoothed H3K36me3 and H3K27me3 histone modification signals within *SOX10* gene body were further plotted using the “loess” function.

## RESULTS

### Design of CellMATE to capture synergistic properties in paired multimodal data

CellMATE is an advanced deep generative framework specifically designed for the early and nonlinear integration of paired sc-multimodal data. Central to this framework is a multi-head adversarial training module (Figure 1), which excels in auto-learning the true distribution and noise characteristics across diverse modalities and integrating these varied attributes into a unified variational autoencoder (VAE) without relying on any predetermined assumptions. The overall architecture allows for the simultaneous utilization of all features, effectively capturing both the additive and synergistic strength in paired sc-multimodal data.

**Figure 1.**
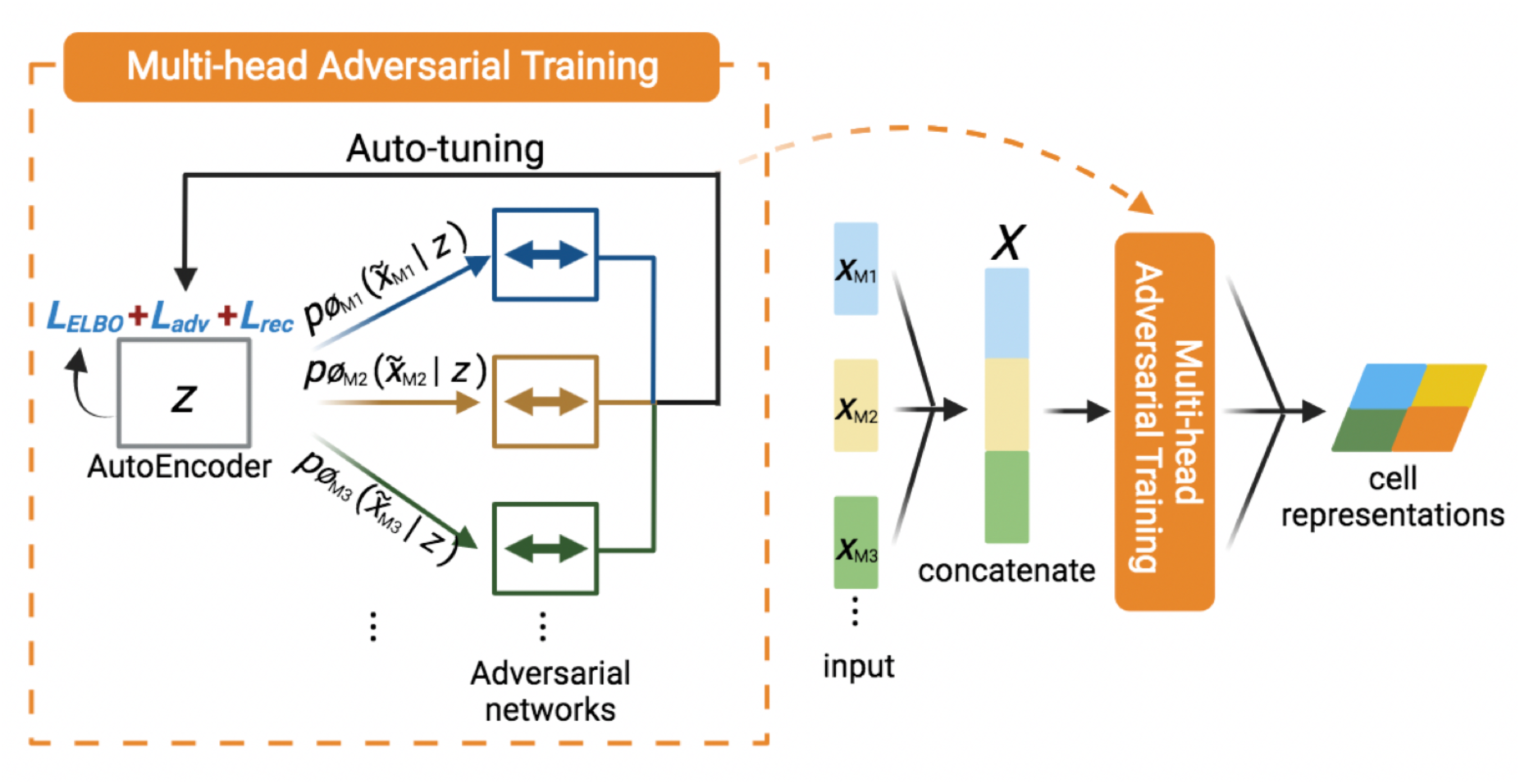
Overview of CellMATE. CellMATE uniquely utilizes a multi-head adversarial training module to enable nonlinear early-integration of sc-multiomics. The input multimodal data (*X*), concatenation of features from all modalities, is simultaneously used to learn a modal-free low-dimensional stochastic latent space *z*. The modal-specific adversarial networks which learn the modal-specific distribution and noise can then auto-tune *z*. Adversarial loss (*Ladv*) and recovery loss (*Lrec*) are introduced alongside the evidence lower bound (*LELBO*) to ensure the accuracy and reliability of *z* and the reconstructed multimodal data.

In operation, multimodal data (*X*), comprising concatenated features from all modalities, is fed into a modal-free encoder to transform the multimodal inputs into a unified, low-dimensional latent space (*z*). This space comprehensively represents a cell by simultaneously utilizing all available features. Consequently, beyond the additive effects as seen in late integration, CellMATE’s ultimate modal-free latent space (*z*) can also capture the cross-modal synergies (Figure 1). To ensure the fidelity of the latent representation, the encoded information is then projected back into the original high-dimensional space, examined and adjusted through modal-specific adversarial networks. These networks ensure the precise recovery of true data distribution and noise for each modality, releasing CellMATE from constraint of modality specificity or dependency on parametric distribution modeling. CellMATE further enhances the training process by incorporating an adversarial loss (*Ladv*) and a recovery loss (*Lrec*), alongside the original evidence lower bound (*Lelbo*). This trio of metrics guides the optimization of the training’s gradient, enhancing the accuracy and reliability of both the latent representations and the reconstructed multimodal data. By leveraging the cross-modal interplay from all modalities, CellMATE captures synergistic advantages of joint profiling, enhances precision of latent representation, and delineates dynamic trajectories.

### CellMATE uniquely harnesses both the additive and synergistic strength of joint profiling

To evaluate the capability of CellMATE to harness the full cross-modal potential of sc-multimodal data, we first applied it on a dataset where the strengths of modalities are uneven. When integrating scRNA with paired low-quality scATAC data[29], CellMATE not only retained the potential of scRNA but also enhanced the performance of the paired data, whereas WNN deteriorated the performance (Figure S1 and S2). Remarkably, CellMATE uniquely retained identification of a grouping of active proliferating cells (Figure S3), a subtle yet significant strength of the RNA modality[29]. These results suggest that CellMATE can better retain the benefit of each modality to enhance the additive strength of joint profiling.

In addition to additive benefit, the early integration design of CellMATE enables the capture of synergistic advantages. In a joint profiling of three histone modifications during human embryonic stem cells (hESCs) differentiation[27], the addition of another modality consistently improved CellMATE’s performance , a benefit not observed with WNN or MOFA+ (Figure 2 A-B). This consistent enhancement was verified through multiple iterations of clustering with random seeds, as evidenced by positive accuracy variance values (Figure 2 C-D). Significantly, CellMATE distinguished hESCs and all the three induced cell lineages, with a particular proficiency in separating ectoderm cells from hESCs (Figure 2E). This unique advantage of CellMATE on joint profiling to separate ectoderm cells is unachievable by any individual modality (Figure 2E), demonstrating CellMATE’s capability in capturing the synergistic benefit in joint profiling.

**Figure 2.**
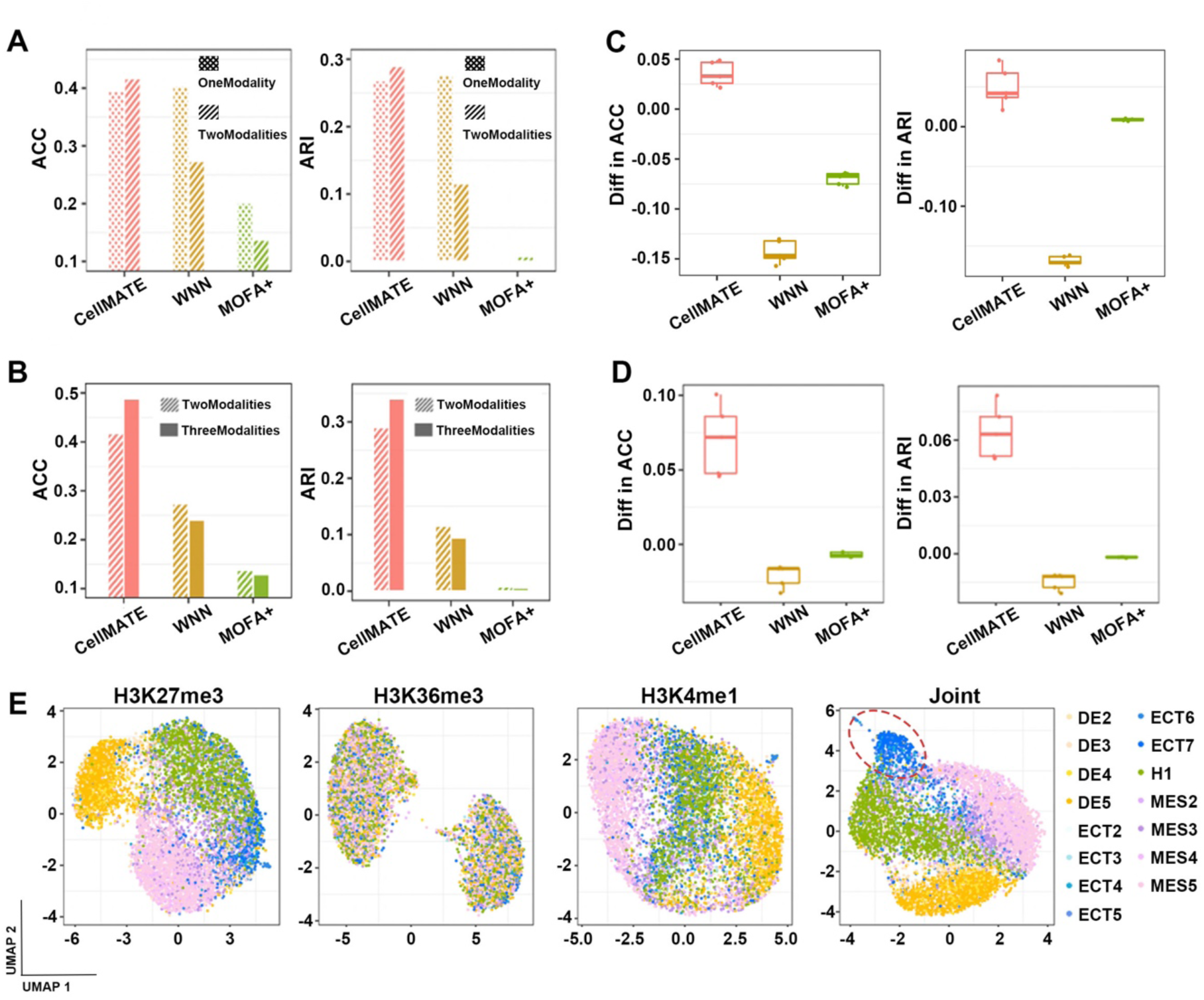
CellMATE captures synergistic advantages of joint profiling. (**A-B**) Bar plots of the clustering performances of CellMATE, WNN and MOFA+ using one or two modalities (**A**) or two or three modalities (**B**), evaluated by unsupervised clustering accuracy (ACC) (left) or Adjusted Rand Index (ARI) (right). (**C-D**) Box plots summarizing the differences between performance for each method with two modalities vs one modality (**C**) or three modalities vs two modalities (**D**). Replicates were performed in 5 runs with different random seeds in the clustering step. Box plots depict the median, quartiles and range. n = 5 for each box. (**E**) UMAP embeddings of the MulTI-Tag dataset using individual modality scChIP (H3K27me3), scChIP (H3K36me3), scChIP (H3K4me1), or joint modalities. Each dot represents a cell colored by cell lineage labels. The ectoderm cells that can only be separated by joint profiling are highlighted.

These findings demonstrate that CellMATE can enhance the additive benefit under modal discrepancy. More crucially, CellMATE can extract the synergistic potential of paired sc-multimodal data. All these results underscore CellMATE’s leading edge in unlocking the full potential of joint profiling.

### CellMATE enhances detection of dynamic plasticity in joint profiling of RNA+ATAC

To evaluate proficiency of CellMATE in inferring transient and delicate cellular states by leveraging the full potential of paired sc-multimodal data, we applied it to the widely used joint profiling like scRNA+scATAC across two platforms, sci-CAR and 10X. The results revealed that CellMATE adeptly identified all cell types within the kidney (sci-CAR)[29] and PBMCs (10X)[32] datasets (Figure S4-S5). We then benchmarked CellMATE against seven state-of-the-art methods for paired sc-multimodal data analysis, including WNN[9], MOFA+[11] , scAI[10], Cobolt[13], MultiVI[14], scMM[12] and scDEC[15]. We quantitatively assessed performance using ARI and ACC metrics, evaluating the congruence between the clusters derived from latent representation and ground truth cell-type labels. CellMATE achieved the highest ARI and ACC scores, outperforming all compared methods in both datasets (Figure 3 A-B).

**Figure 3.**
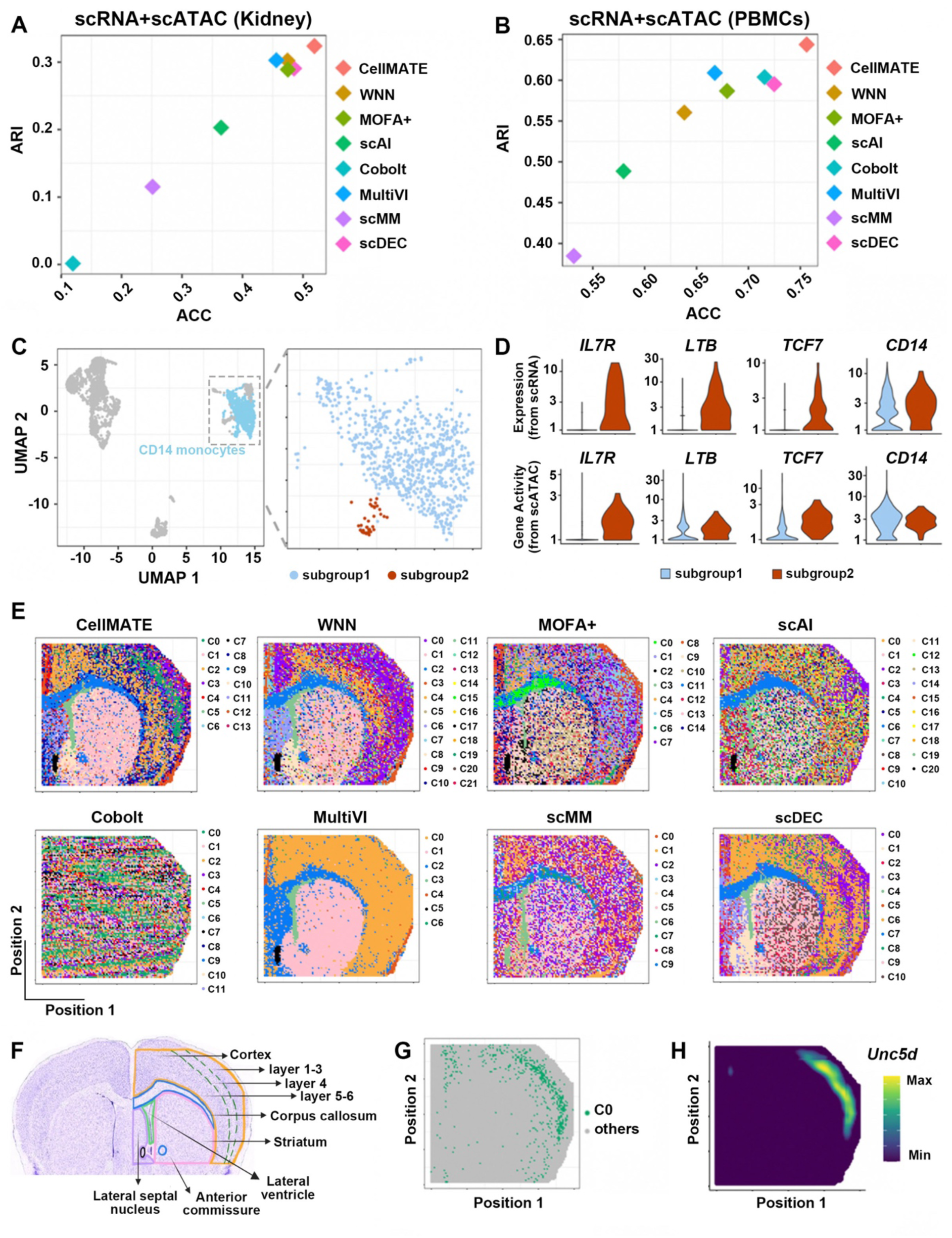
CellMATE uniquely unveils cellular plasticity and nuanced spatial structure on joint profiling of transcriptome and chromatin accessibility. (**A-B**) Clustering performance of CellMATE and the competing methods evaluated by ACC (X axis) and ARI (Y axis) on scRNA+scATAC datasets profiled on sci-CAR kidney (**A**) and 10X PBMCs (**B**). (**C)** UMAP embeddings of the 10X PBMCs dataset learned by CellMATE, with CD14 monocytes highlighted (left). The inset shows the subgroups of CD14 monocytes uncovered by CellMATE (right). (**D**) Expression from scRNA (top) and gene activity from scATAC (bottom) of marker genes of stimulated monocytes in the two subgroups of CD14 monocytes. (**E**) Spatial distribution of all clusters learned by CellMATE or competing methods in spatial RNA+ATAC of mouse brain. Each dot is colored by cluster labels. (**F**) Main structures in the mouse brain. (**G**) Spatial distribution of cluster0 (C0) from CellMATE in spatial RNA+ATAC of mouse brain. (**H**) Spatial mapping accessibility of *Unc5d*, a marker gene of layer 4 of cortex, in spatial RNA+ATAC.

The enhanced latent inference generated by CellMATE enables the discernment of subtle dynamic cell states. For instance, within the PBMC dataset, CellMATE identified distinct subgroups of CD14 monocytes (Figure 3C and Figure S6). Although both subgroups exhibited active features at *CD14*, subgroup2 was characterized by elevated transcriptional and accessible features at *IL7R*, *LTB* and *TCF7* (Figure 3D), which were characteristics of stimulated monocytes[33].

In addition to scRNA+scATAC technology, joint profiling of spatial transcriptome and chromatin accessibility represents a cutting-edge advance[34, 35]. We further applied CellMATE to spatial paired data of mouse brain[36]. CellMATE, alongside WNN, identified all major and finer brain structures, including cortex, corpus callosum, striatum, lateral ventricle and lateral septal nucleus (Figure 3 E-F and Figure S7). Notably, CellMATE uniquely distinguished additional cortical layers. The layer C0 revealed by CellMATE was marked by high accessibility at *Unc5d* (Figure 3 G-H), which is a characteristic gene of layer 4 in the cortex[25]. This layer was undetectable using any other methods (Figure S7). These outcomes demonstrate that CellMATE generates superior latent representations of RNA+ATAC joint profiling, which facilitates detection of dynamic cellular plasticity and spatial structure.

### CellMATE is robust and effective in diverse joint profiling scenarios

Recent advancements in experimental techniques have expanded the repertoire and scale of measurable modalities within single cells beyond the scRNA+scATAC combination. However, computational methods optimized for specific multimodal combination may falter when applied to others due to the distinct distribution and noise attribute of each modality. To ascertain CellMATE’s adaptability, we conducted an extensive investigation of its performance in various dual-modal datasets, as well as triple-modal datasets.

This investigation of dual-modal datasets encompassed joint profiling of single cell Antibody-Derived Tags (scADT) and scATAC (ASAP-seq)[23], scADT and scRNA (CITE-seq)[23], scChIP of different histone modifications (MulTI-Tag)[27], and scATAC and scChIP (nanobody-based scCUT&Tag)[26]. The results showed that CellMATE adeptly inferred latent representations and successfully elucidated cellular heterogeneity across all datasets (Figure 4A and Figure S8-S11). The benchmarking revealed that CellMATE’s performance was favourable or comparable across all these modality combinations, as assessed using ARI and ACC metrics (Figure 4B), further supported by the local inverse Simpson’s index (LISI) and overcorrection (OC) score (Figure S12).

**Figure 4.**
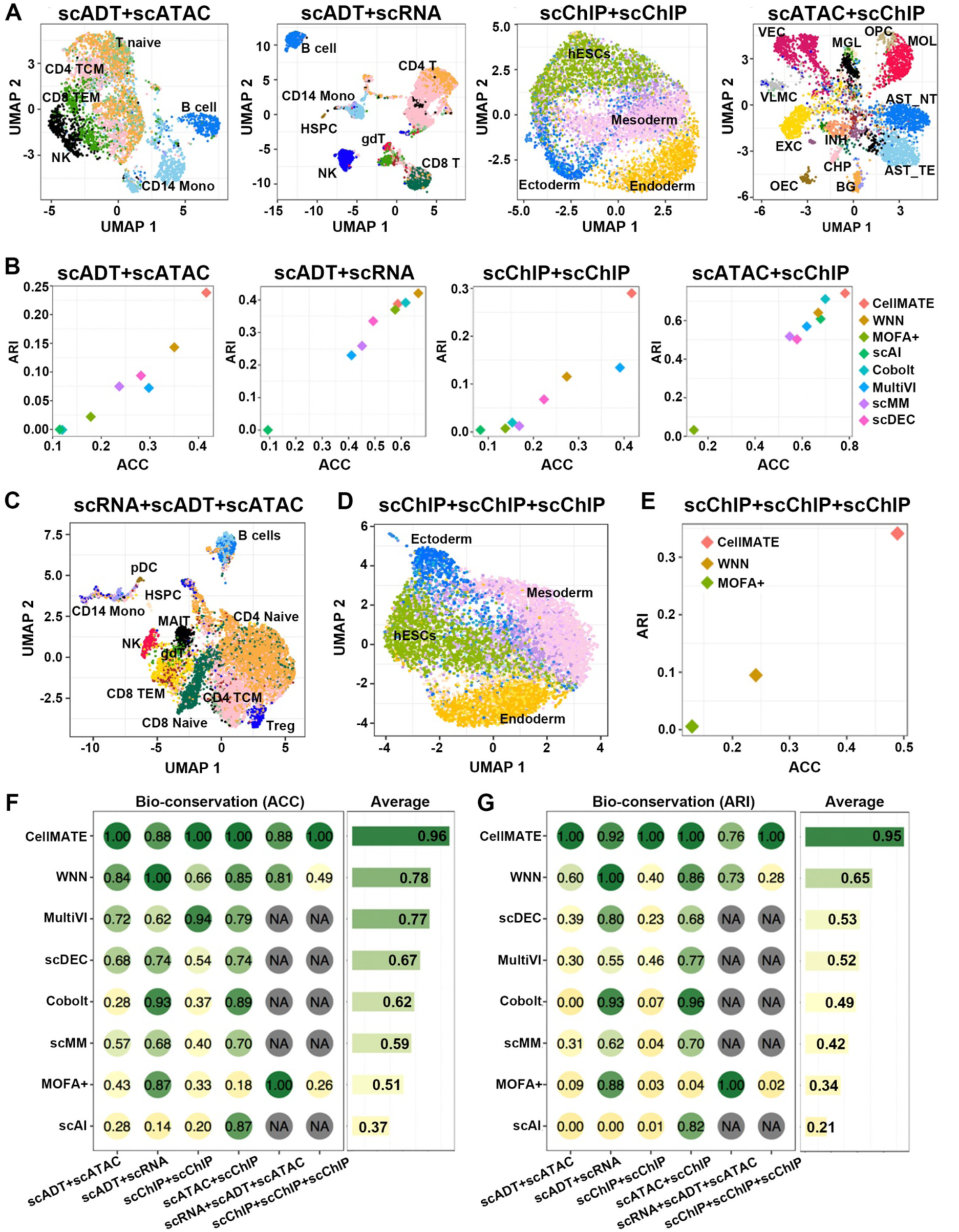
CellMATE is robust on joint profiling with different combinations and numbers of modalities. (**A**) UMAP embedding learned by CellMATE of scADT+scATAC (ASAP-seq), scADT+scRNA (CITE-seq), scChIP+scChIP (MulTI-Tag) and scATAC+scChIP (scCUT&Tag) datasets. Each dot represents a cell colored by cell type labels. (**B**) Clustering performance of CellMATE and the competing methods evaluated by ACC (X axis) and ARI (Y axis) on each dataset. (**C-D**) UMAP embeddings learned by CellMATE of the scRNA+scADT+scATAC (DOGMA-seq) dataset (**C**) and scChIP+scChIP+scChIP (MulTI-Tag) dataset (**D**). Each dot represents a cell colored by cell type or lineage labels. (**E**) Clustering performance of CellMATE, WNN and MOFA+ on the MulTI-Tag dataset, evaluated by ACC (X axis) and ARI (Y axis). (**F-G**) Bubble plots summarizing the performance of the methods against the best performance on each dataset by ACC (**F**) and ARI (**G**). Color represents the relative ACC or ARI score of each method against the best performance score on each dataset. NA stands where the method is not applicable.

Expanding our investigation to more modalities within a single cell, CellMATE’s proficiency was also tested in sc-triple-modal scenarios, such as scRNA+scADT+scATAC (DOGMA-seq)[23] and scChIP of three distinct histone modifications (MulTI-Tag)[27]. CellMATE successfully identified cell types in the DOGMA-seq PBMCs dataset, including Treg and HSPC cells (Figure 4C and Figure S13). More impressively, CellMATE showcased its unique strength in deciphering the dynamic processes like differentiation. On the MulTI-Tag dataset, only CellMATE distinguished hESCs and all the three induced cell lineages (Figure 4D and Figure S14), achieving the highest ARI and ACC scores (Figure 4E). CellMATE significantly outperformed other methods capable of analyzing sc-triple-modal data, such as WNN and MOFA+.

We evaluated the bio-conservation quality of cell embeddings generated by each method. By comparing their performance against the best performance on each dataset, CellMATE outperformed the other seven methods examined, ranking highest based on the average scores across the datasets, followed by WNN (Figure 4 F-G). On average, CellMATE achieved ACC bio-conservation scores ranging between 0.88 and 1.00. In contrast, methods like MOFA+ was highly sensitivity to the modalities, with its ACC bio-conservation scores plummeting from 1.00 to 0.18 when RNA modality is not available.

The comprehensive evaluation underscores CellMATE’s exceptional ability to navigate and interpret the intricacies of sc-multimodal data across various scenarios, establishing it as a versatile and reliable tool for leveraging the power of paired sc-multimodal data.

### CellMATE facilitates unravelling the dynamic landscape of differentiation

With the leveraged latent inference from CellMATE, we explored the differentiation trajectories of hESC in MulTI-Tag dataset. CellMATE elucidated all three differentiation trajectories consistent with experimental design, including a previously unreported trajectory from hESCs to ectoderm (Figure 5A). This discovery allowed for a more comprehensive mapping of lineage paths, facilitating the identification of pivotal features along these differential routes (Figure 5B and Figure S15). We found key regulatory genes on the ectoderm path, such as *SOX10*, a core ectoderm cell identity regulator[37] (Figure 5B). The analysis of paired sc-multimodal data further disclosed dynamic histone modifications accompanying ectoderm differentiation. Specifically, it was observed that a reduction in H3K27me3 occurred prior to an increase in H3K36me3 at the *SOX10* locus as hESCs progressed towards the ectodermal fate (Figure 5C). The capability of CellMATE to harness both the additive and synergistic advantages, makes it a standout tool for comprehensively elucidating dynamic biological processes from paired sc-multimodal data.

**Figure 5.**
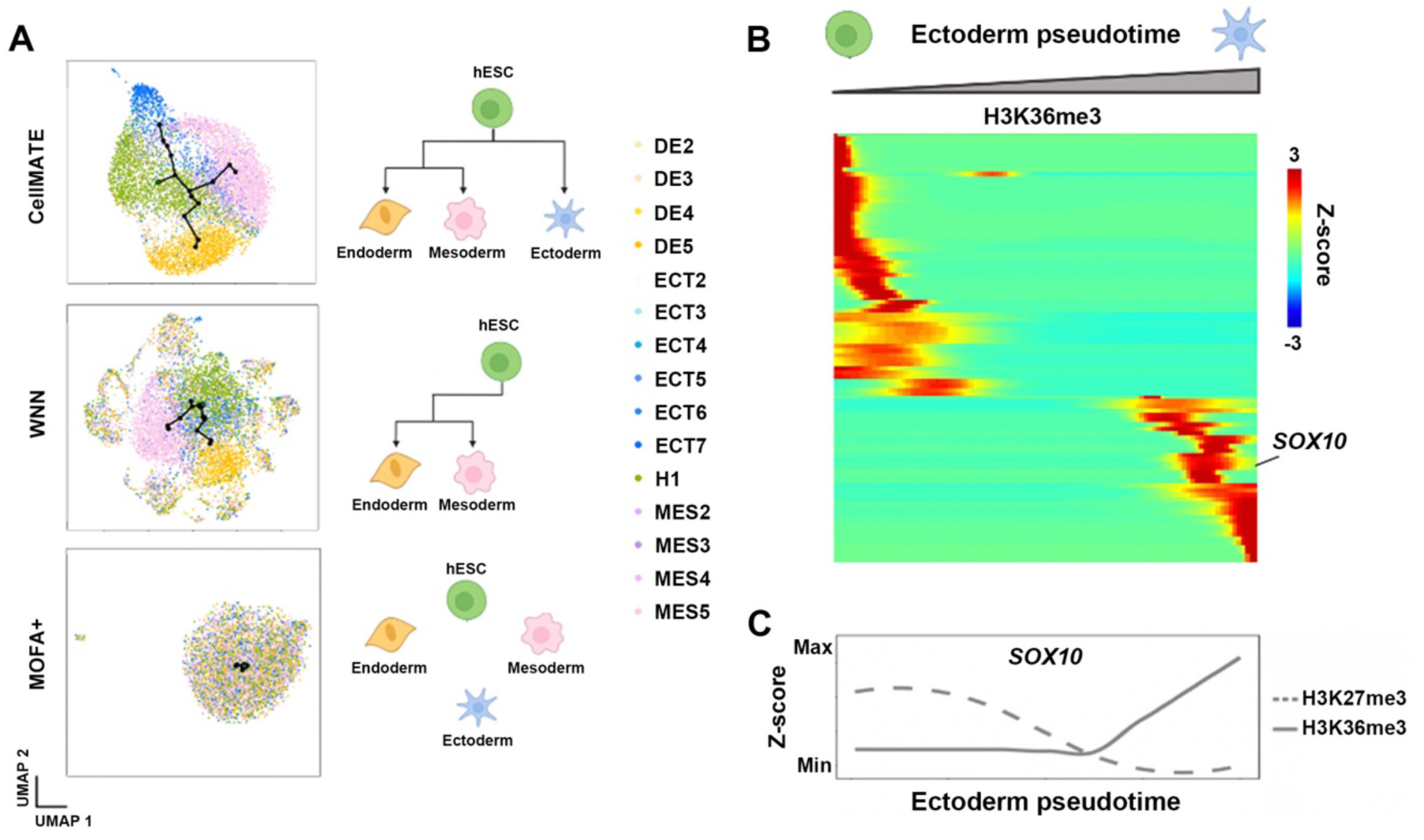
The synergistic interplay detected by CellMATE elucidates dynamic paths during differentiation. (**A**) UMAP embedding learned by CellMATE, WNN or MOFA+ on MulTI-Tag dataset (left) and cartoon diagrams of paths revealed based on cellular representation from each method (right). Each dot represents a cell colored by cell lineage label. Computationally derived trajectories learned by Slingshot are denoted by solid lines. (**B**) Heatmap for top 100 genes (indicated by H3K36me3 measurement in gene bodies) significantly changed along the differentiation trajectory from hESC to ectoderm. Color represents the z-score of normalized H3K36me3 signal. (**C**) Z-score of normalized H3K36me3 and H3K27me3 signal at *SOX10* gene body regions along the differentiation from hESC to ectoderm.

Overall, CellMATE offers a comprehensive cell representation by harnessing the full advantages of paired sc-multimodal data. It provides deeper insights into cell heterogeneity and plasticity, and also provides an enhanced resolution for dissecting the cross-modal interactions.

## DISCUSSION AND CONCLUSION

In this study, we introduced CellMATE, an advanced deep generative model designed for early and non-linear integration of paired sc-multimodal data. CellMATE is unique to capture the synergistic advantages of paired sc-multimodal data, and harness the full potential of joint profiling. CellMATE thus can provide novel biological understanding into dynamic plasticity of cells in addition to steady-state heterogeneity.

The key strength of CellMATE lies in its ability to leverage the full potential of various paired sc-multimodal data. Unlike late integration methods, which operate under the assumption that modalities do not define conflicting sets of cell states[9], an assumption might be violated, CellMATE’s framework is adept at incorporating distinct modal strengths without presupposing their concordance. This is crucial in instances where modalities diverge, such as histone modification priming preceding gene expression changes[38], or where histone modification dynamics signal cellular differentiation, like the loss of H3K27me3 followed by an increase in H3K36me3 at *SOX10* during ectoderm differentiation. CellMATE effectively prevents such discrepancies from compromising the strength of paired sc-multimodal data, a potential pitfall of late integration strategies. Instead, CellMATE takes advantage of these discrepancies to facilitate uncovering of unexpected insights in dynamic processes.

Furthermore, while existing deep generative models like totalVI[21] have optimized the integration of specific modalities, such as protein and RNA profiles, these models falter when confronted with additional or different modalities. CellMATE is empowered by adversarial training, automatically captures the true distribution and noise of each modality. Its capacity to integrate an unrestricted types and numbers of modalities without relying on predetermined parametric modeling positions CellMATE as a versatile tool that can keep pace with the advancing joint profiling experimental techniques.

In conclusion, CellMATE represents a significant step forward in exploring the joint profiling. Its capabilities to harness both the additive and synergistic potential of paired data are particularly valuable in studying dynamic biological processes such as development, differentiation, and diseases.

## DATA AVAILABILITY

All datasets used in this paper were obtained from the original publications and releases. The 10X PBMCs dataset is available from the official website: https://www.10xgenomics.com/resources/datasets. The sci-CAR dataset was downloaded from GEO, accession number GSM3271044. The spatial RNA+ATAC dataset was obtained from the UCSC Cell and Genome Browser: https://brain-spatial-omics.cells.ucsc.edu. The MulTI-Tag dataset was downloaded from the Zenodo repository: https://doi.org/10.5281/zenodo.6636675. The scCUT&Tag dataset was downloaded from GEO, accession number GSE198467. The ASAP-seq, CITE-seq and DOGMA-seq datasets were downloaded from GEO, accession number GSE156478.

## CODE AVAILABILITY

CellMATE is implemented in python and is publicly available via Github (https://github.com/labYangNJU/CellMATE). For reproducibility, code scripts used in this paper are publicly accessible on Github (https://github.com/labYangNJU/CellMATE/tree/main/Reproducibility).

