## Supplementary Figures for "Unlocking cross-modal interplay of single-cell and spatial joint profiling with CellMATE"

**Figure S1.** UMAP embeddings of scATAC-seq from the kidney dataset learned by CellMATE, WNN and MOFA+.

**Figure S2.** The performances of methods using scATAC-only, scRNA-only, or joint analysis on the kidney dataset evaluated by ACC.

**Figure S3.** Matrices of the proportions of active proliferating cells in each cluster learned by different methods.

**Figure S4.** UMAP embeddings of the kidney dataset learned by CellMATE or competing methods.

**Figure S5.** UMAP embeddings of the 10X PBMCs dataset learned by CellMATE or competing methods.

**Figure S6.** Subgroup2 of CD14 monocytes from 10X PBMCs dataset in CellMATE or competing methods.

**Figure S7.** Spatial distribution of all clusters from CellMATE or competing methods in spatial RNA+ATAC of mouse brain.

**Figure S8.** UMAP embeddings of the ASAP-seq dataset learned by CellMATE or competing methods.

**Figure S9.** UMAP embeddings of the CITE-seq dataset learned by CellMATE or competing methods.

**Figure S10.** UMAP embeddings of the MulTI-Tag dataset (scChIP H3K27me3 + scChIP H3K36me3) learned by CellMATE or competing methods.

**Figure S11.** UMAP embeddings of the nanobody-based scCUT&Tag dataset learned by CellMATE or competing methods.

**Figure S12.** Assessment of CellMATE and the competing methods on cell-type separation.

**Figure S13.** UMAP embeddings of the DOGMA-seq dataset learned by CellMATE, WNN and MOFA+.

**Figure S14.** UMAP embeddings of the MulTI-Tag dataset (scChIP H3K27me3 + scChIP H3K36me3 + scChIP H3K4me1) learned by CellMATE, WNN and MOFA+.

**Figure S15.** Heatmap for top 100 peaks of H3K27me3 (left) or H3K4me1 (right) significantly changed along the differentiation trajectory from hESC to ectoderm.

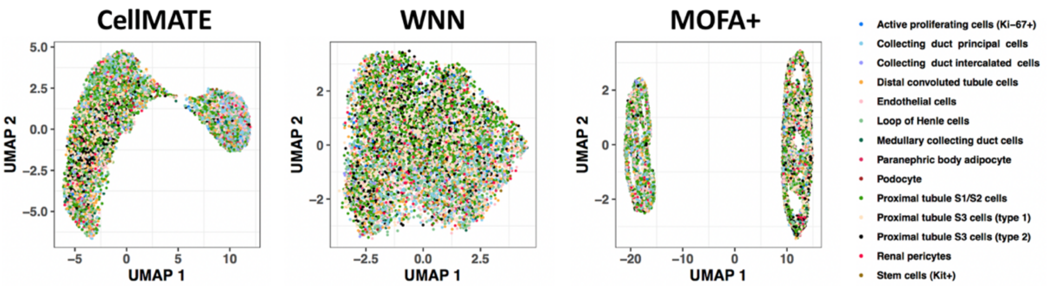


**Figure S1. UMAP embeddings of scATAC-seq from the kidney dataset learned by CellMATE, WNN and MOFA+.** Each dot represents a cell colored by cell type labels.

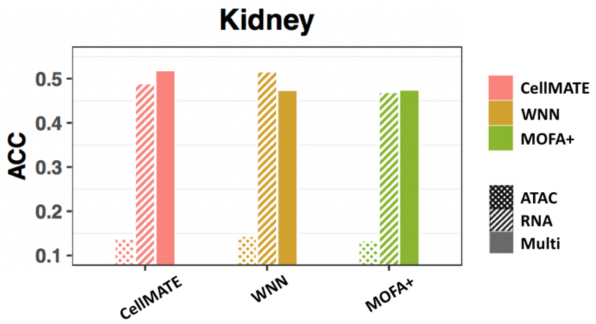


**Figure S2. The performances of methods using scATAC-only, scRNA-only, or joint analysis on the kidney dataset evaluated by ACC.**

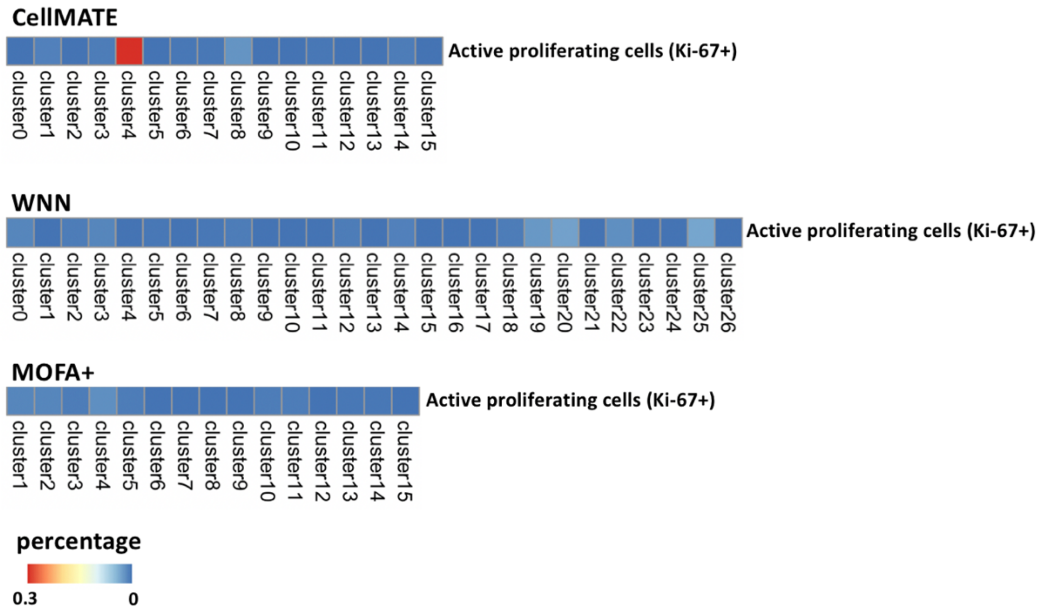


**Figure S3. Matrices of the proportions of active proliferating cells in each cluster learned by different methods.** Color represents the percentage of active proliferating cells in each cluster.

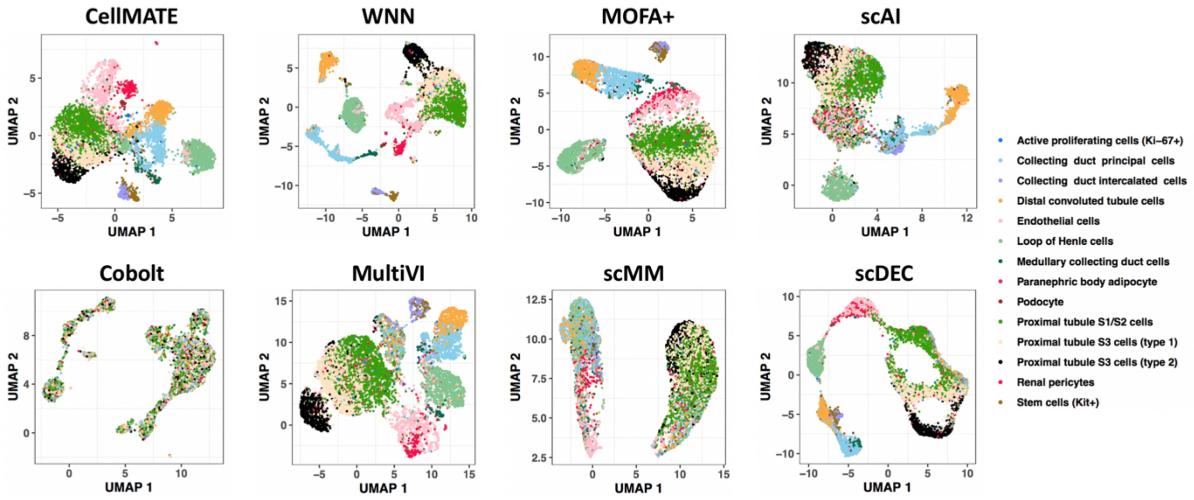


**Figure S4. UMAP embeddings of the kidney dataset learned by CellMATE or competing methods.** Each dot represents a cell colored by cell type labels.

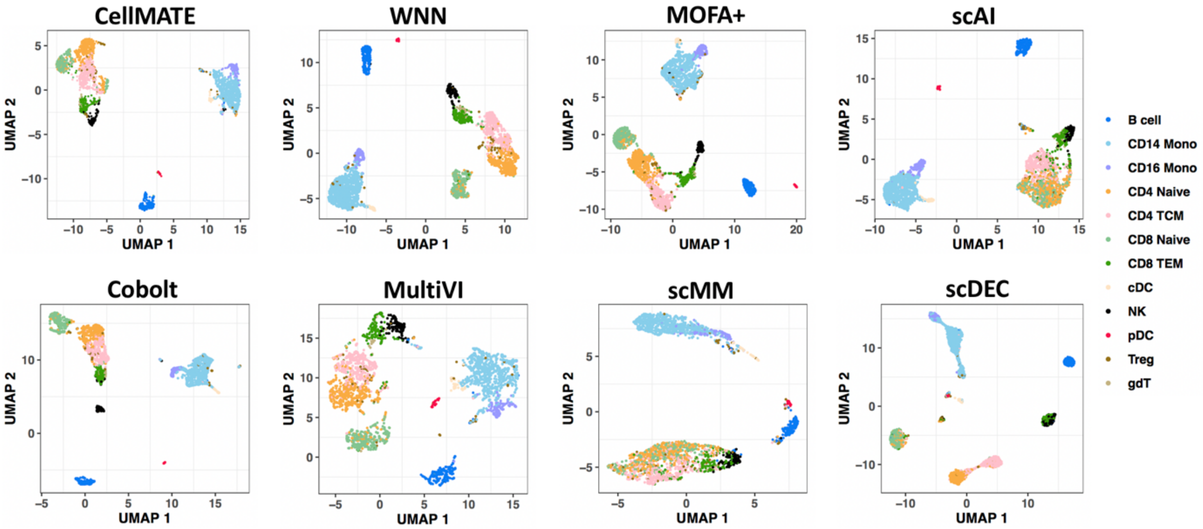


**Figure S5. UMAP embeddings of the 10X PBMCs dataset learned by CellMATE or competing methods.** Each dot represents a cell colored by cell type labels. Cell types with more than 20 cells are shown.

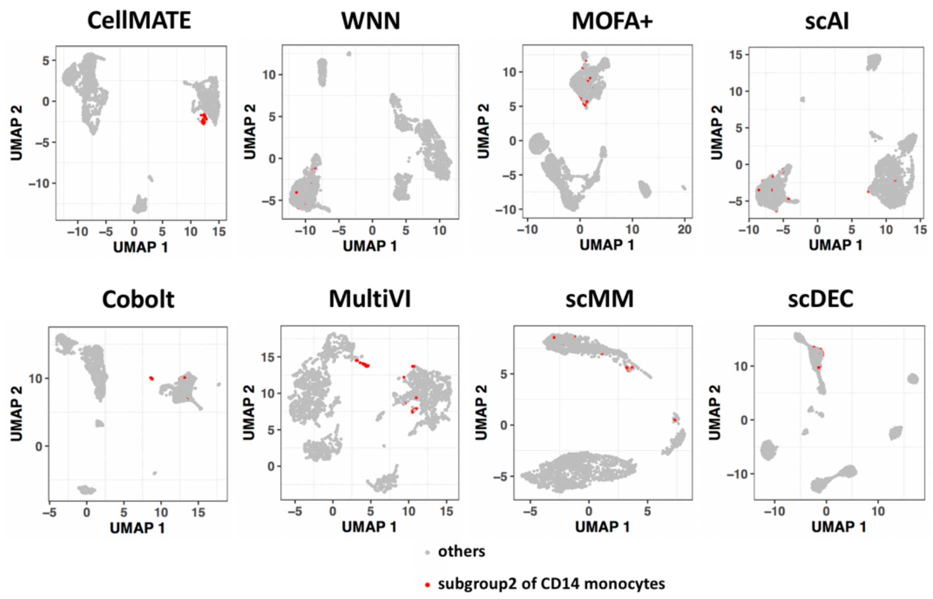


**Figure S6. Subgroup2 of CD14 monocytes from 10X PBMCs dataset in CellMATE or competing methods.** Subgroup2 of CD14 monocytes are highlighted.

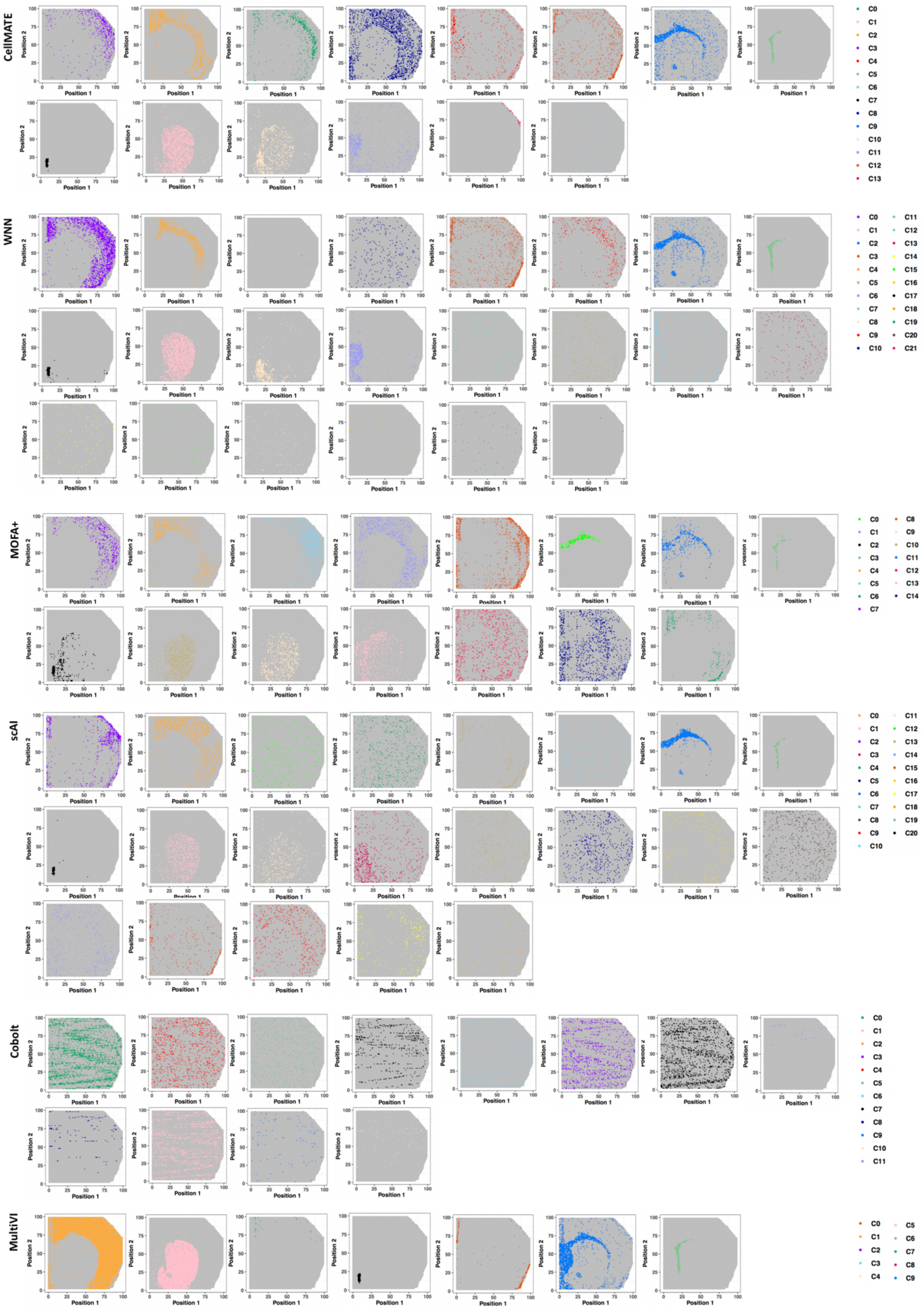
CellMATE
WNN
MOFA+
scAI
Cobolt
MultiVI
Position 1
Position 2
C0 C1 C2 C3 C4 C5 C6 C7 C8 C9 C10 C11 C12 C13
C0 C1 C2 C3 C4 C5 C6 C7 C8 C9 C10 C11 C12 C13 C14 C15 C16 C17 C18 C19 C20 C21
C0 C1 C2 C3 C4 C5 C6 C7 C8 C9 C10 C11 C12 C13 C14
C0 C1 C2 C3 C4 C5 C6 C7 C8 C9 C10 C11 C12 C13 C14 C15 C16 C17 C18 C19 C20
C0 C1 C2 C3 C4 C5 C6 C7 C8 C9 C10 C11
C0 C1 C2 C3 C4 C5 C6 C7 C8 C9

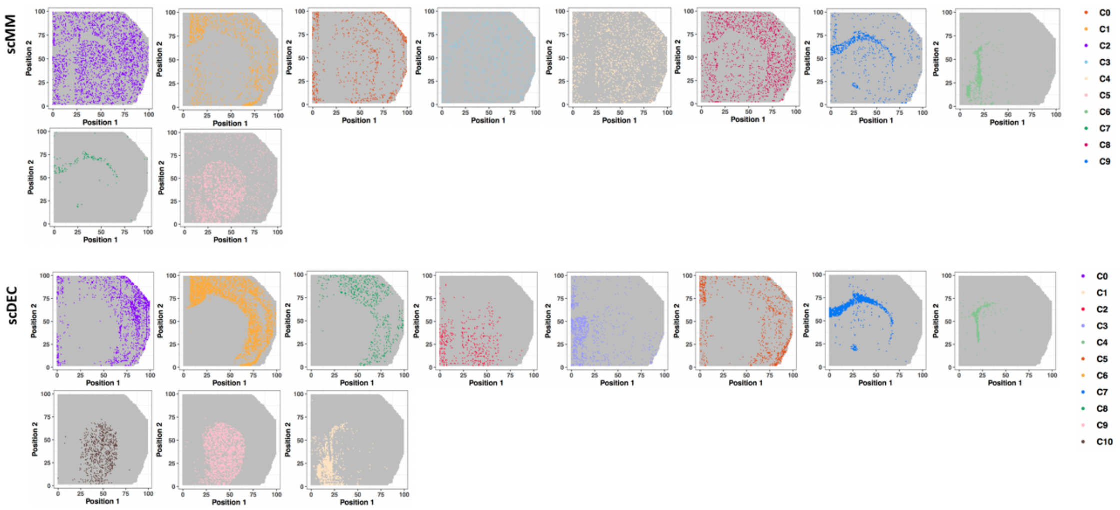


**Figure S7. Spatial distribution of all clusters from CellMATE or competing methods in spatial RNA+ATAC of mouse brain.** Each dot represents a cell colored by cluster labels.

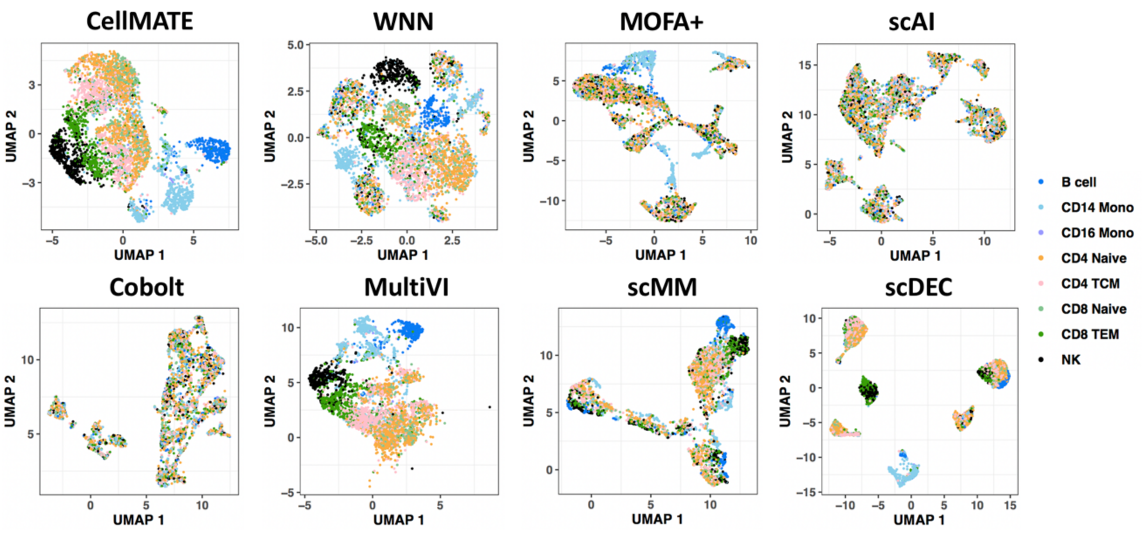


**Figure S8. UMAP embeddings of the ASAP-seq dataset learned by CellMATE or competing methods.** Each dot represents a cell colored by cell type labels. Cell types with more than 20 cells are shown.

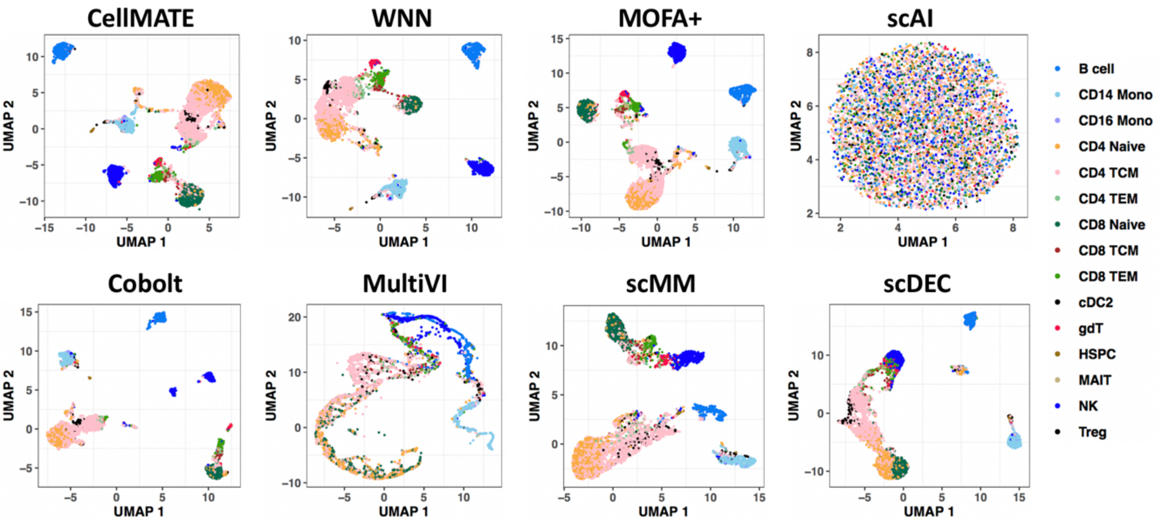


**Figure S9. UMAP embeddings of the CITE-seq dataset learned by CellMATE or competing methods.** Each dot represents a cell colored by cell type labels. Cell types with more than 20 cells are shown.

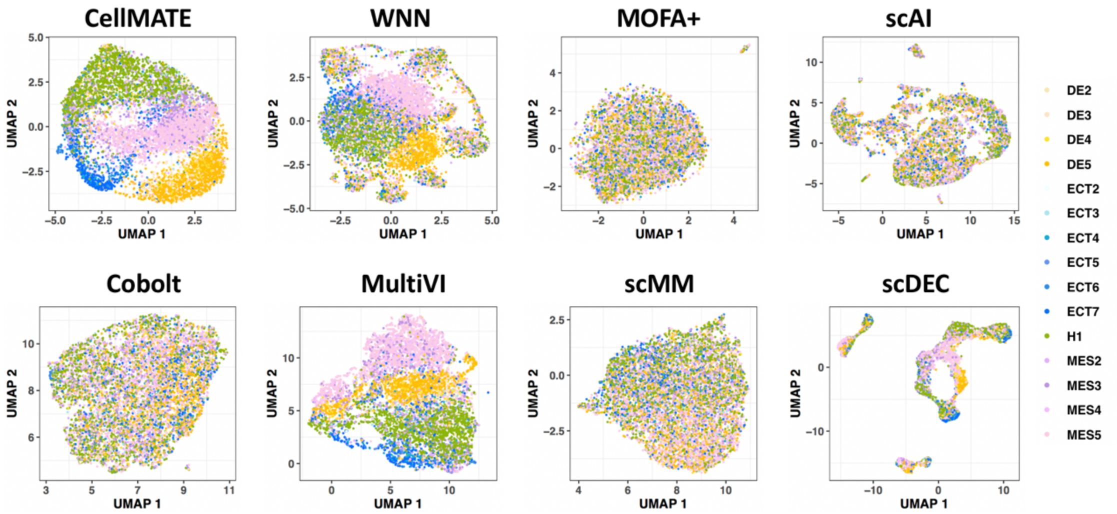


**Figure S10. UMAP embeddings of the MulTI-Tag dataset (scChIP H3K27me3 + scChIP H3K36me3) learned by CellMATE or competing methods.** Each dot represents a cell colored by cell lineage labels.

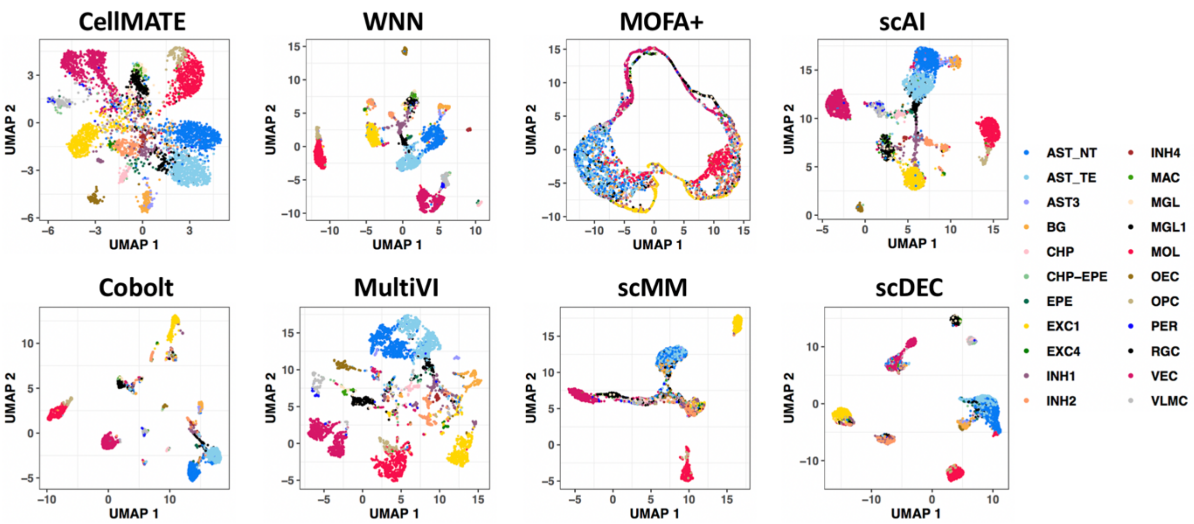


**Figure S11. UMAP embeddings of the nanobody-based scCUT&Tag dataset learned by CellMATE or competing methods.** Each dot represents a cell colored by cell type labels. Cell types with more than 20 cells are shown.

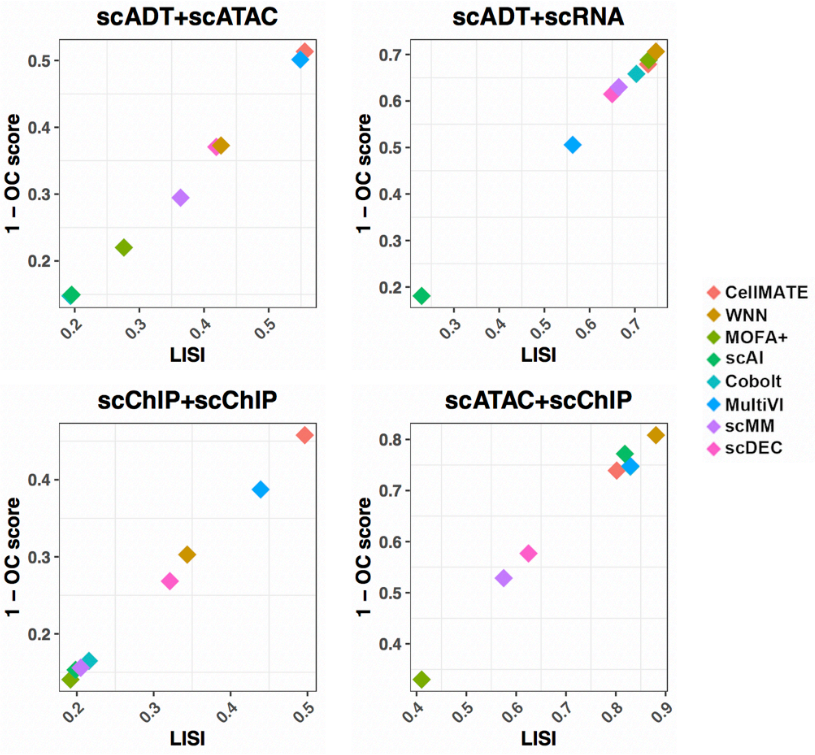


**Figure S12. Assessment of CellMATE and the competing methods on cell-type separation.** Cell-type separation performance of CellMATE and the competing methods evaluated by LISI (X axis) and (1 – OC) score (Y axis) on each dataset.

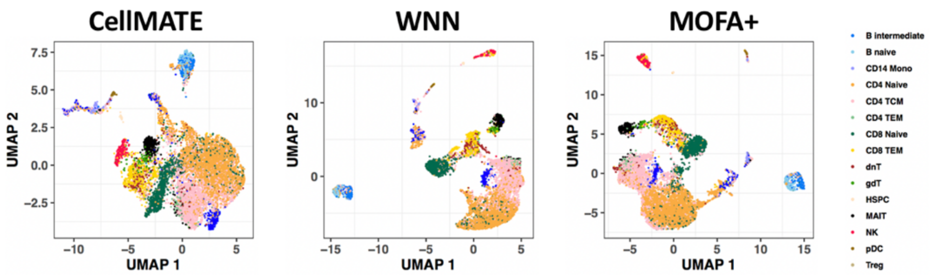


**Figure S13. UMAP embeddings of the DOGMA-seq dataset learned by CellMATE, WNN and MOFA+.** Each dot represents a cell colored by cell type labels. Cell types with more than 20 cells are shown.

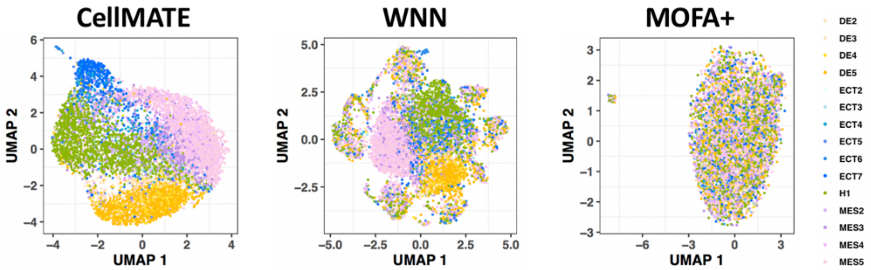


**Figure S14. UMAP embeddings of the MulTI-Tag dataset (scChIP H3K27me3 + scChIP H3K36me3 + scChIP H3K4me1) learned by CellMATE, WNN and MOFA+.** Each dot represents a cell colored by cell lineage labels.

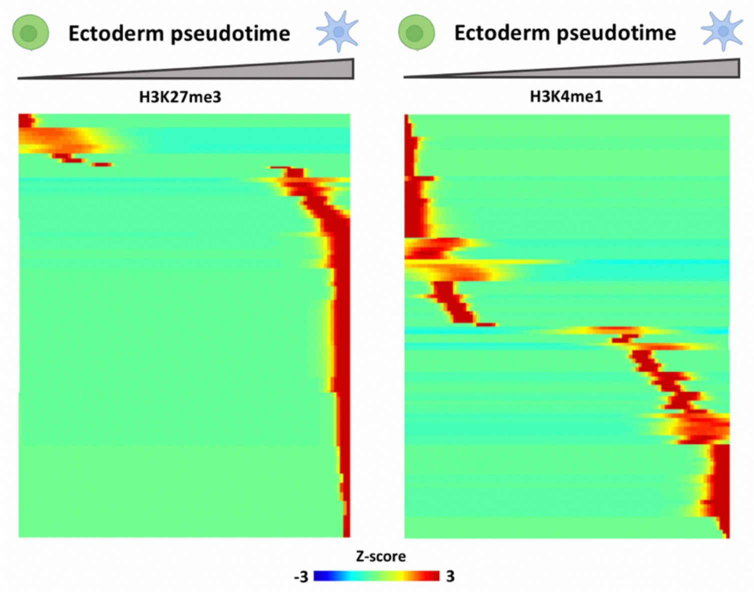


**Figure S15. Heatmap for top 100 peaks of H3K27me3 (left) or H3K4me1 (right) significantly changed along the differentiation trajectory from hESC to ectoderm.** Color represents the z-score of normalized H3K27me3 or H3K4me1 signal.
